# Evaluation of Postharvest Storability of Ponkan Mandarins at Different Storage Temperatures Based on Principal Component Analysis

**DOI:** 10.1101/841379

**Authors:** Nan Cai, Chunpeng Wan, Jinyin Chen, Chuying Chen

**Affiliations:** Jiangxi Key Laboratory for Postharvest Technology and Non-destructive Testing of Fruits & Vegetables, Collaborative Innovation Center of Post-harvest Key Technology and Quality Safety of Fruits and Vegetables in Jiangxi Province, Jiangxi Agricultural University, Nanchang 330045, China; (N.C.); (C.W.); Pingxiang University, Pingxiang 337055, China

**Author notes:** Correspondence (J.C.); (C.C.); (J.C.&C.C.).

**Keywords:** Ponkan mandarins, Storage temperature, Preservation storability, Principal component analysis

## Abstract

To reduce postharvest losses of Ponkan mandarins caused by outdated storage facilities and preservation technology, we evaluated the preservation effect of different storage temperatures on Ponkan mandarins (5 ±1, 10 ± 1, 15 ± 1, and 20 ±1 °C), and obtained a comprehensive score using principal component analysis (PCA) to determine its suitable storage temperature. The results indicate that, relative to the other three storage temperatures, storage at 10 °C significantly maintains high total soluble solid content, titratable acid, and vitamin C contents; the accumulation of malondialdehyde (MDA) and hydrogen peroxide (H_2_O_2_) content decreased and changes in the relative conductivity (REC) were suppressed; and high activities of superoxide-dismutase (SOD), peroxidase (POD), catalase (CAT), and ascorbate peroxidase (APX), as well as high contents of total phenol and total flavonoid were maintained. The PCA and clustering heat map results show that that the comprehensive score was the highest when stored at 10 °C. The data indicate that the suitable storage temperature of Ponkan mandarins at 10 °C significantly decreased MDA accumulation and reactive oxygen species metabolism, maintains high antioxidant capacity, maintains good fruit quality and achieves good storage and preservation effect, which is the appropriate storage temperature for Ponkan mandarins.

## Introduction

Ponkan mandarins (*Citrus reticulata Blanco*, PM) are one of the main broad-skin citrus cultivars and are widely cultivated in Southern China. Jing’an Ponkan mandarin is a famous mandarin variety in Jiangxi Province, characterized by a compact plant type, early fruit bearing, rounded fruit shape, bright color, sweet taste, crisp flesh, storage resistance, and late maturity. These mandarins are a major source of economic income for fruit growers in the Jing’an area [1].

Appropriate postharvest treatment can significantly reduce fruit loss, improve fruit quality, and result in higher profits. However, citrus fruit may exhibit various disorders during harvest that limit the storage period and reduce their commercial value [2]. Many factors affect the postharvest storage of fresh fruit, such as low temperature storage, mechanical damage by harvesting and transporting, and harvest periods [3–5]. The most common problem during the postharvest storage of citrus is decay. Many reports indicate that the use of polyethylene film bagging [6], the application of 1-methylcyclopropene [7], and coating with plants extracts or its main antifungal constituents, such as cinnamaldehyde, limonene, and clove essential oil [8], during postharvest treatment can effectively decrease the incidence of decay in citrus fruits during storage.

Storage temperature affects respiration [9], fruit texture [10], and fruit physiological metabolism [11]. During storage, appropriate storage temperature can reduce the occurrence of pests and diseases [12]; decrease the fruit rot rate [13]; maintain the luster and flavor of the fruit [14]; increase soluble solids (TSS) and titratable acid (TA) contents [15]; delay fruit senescence [16]; and increase antioxidant enzyme activity and maintain higher peroxidase (POD), catalase (CAT), superoxide dismutase (SOD), and ascorbate peroxidase (APX) activities [17]. To some extent, the appropriate storage temperature can inhibit the accumulation of MDA content. A too high or too low storage temperature is harmful to fruits. Therefore, determining the suitable storage temperature for Ponkan mandarins after harvest is crucial. Studies have reported that long-term storage of Valencia oranges at high storage temperatures of 20 and 25 °C leads to a decrease in the flavor quality of the oranges and the accumulation of stale flavors [18]. Compared with grapefruit stored at higher (12 °C) or lower (4 °C) temperatures, long-term storage at 8 °C at a medium temperature can better maintain their flavor [19]. Therefore, the minimum safe temperature for postharvest storage of citrus is suggested to be between 5 and 8 °C [20,21]. The susceptibility of citrus fruits to chilling injury varies with cultivars. When stored in the range of 2 to 6 °C, chilling injury may occur in grapefruit, Shamouti orange, Vorschach orange, and lemon fruit [22]. Jose et al. also found that chilling injury reduced the appearance quality and commercial value of citrus fruits [23]. Therefore, to determine the best storage temperature of Ponkan mandarins provides a scientific basis for reducing fruit loss after harvest and guidance for the production and harvest of fruit for farmers.

The suitable storage conditions of fruit cannot be reflected by a single index, but requires a comprehensive analysis of multiple indexes. Principal component analysis (PCA) is an analysis method that transforms multiple variables into multiple comprehensive variables. [24]. Then, the correlation between different variables is explained by Pearson correlation analysis. In this study, we used PCA to establish an evaluation system for the indexes related to treatments at different storage temperatures, which could be used as a basis for comprehensively evaluating Ponkan mandarins storage and preservation under different storage temperatures. The suitable storage temperatures of Ponkan mandarins were screened out, providing a theoretical basis for actual production to reduce losses and thereby help increase profit.

## Materials and Methods

### Experimental Materials and Storage Conditions

Fresh Ponkan mandarins were harvested at 195 days after full-bloom (DAF) stage from a commercial orchard in the Xiangtian district of Jing’an City (Jiangxi Province, China), and transported back to the Jiangxi Provincial Key Laboratory of Fruit and Vegetable Preservation and Nondestructive Testing (Nanchang, Jiangxi, China) within 4 h. After sweating for 3 days, the healthy fruits without diseases or mechanical damage and uniform in color, size, shape, and maturity were selected for use throughout this experiment. Then, the selected fruits were randomly separated into 4 lots, and stored at 5 ± 1 °C, 10 ± 1, 15 ± 1, and 20 ± 1 °C with a relative humidity of 85%–90% for 120 days. We selected 15 fruits every 15 days to evaluate their physiological and biochemical indexes.

### Physiological Analysis of Ponkan Mandarins

#### Decay Rate

The decay rate of Ponkan mandarins due to peel browning and diseases was measured and calculated on the initial fruit number for each lot every 15 days, and is expressed in percentage.

#### Weight Loss

The weight loss of Ponkan mandarins was measured and calculated on the initial fruit weight basis for each lot every 15 days, and is expressed in percentage.

#### Light (L*) and Citrus Color Index (CCI)

The L* and CCI was measured using a MINOLTA CR-400 colorimeter (D65 light source; Konica Minolta Sensing, Inc., Osaka, Japan) for 15 randomly selected fruits, described following the method reported by Chen et al. [25] with slight modifications. Four points were taken near the equator of the fruit for evaluation.

### Biochemical Analysis of Ponkan Mandarins

#### Titratable Acid (TA) and Total Soluble Solid (TSS)

The TA content was determined by titration with 0.1 N sodium hydroxide to pH 8.1, and is expressed as a percentage of citric acid, which is the diagnostic organic acid in citrus fruits.

The TSS content was measured with the aid of a RA-250WE Brix-meter (Atago, Tokyo, Japan), and the result is expressed as a percentage.

#### Determination of Antioxidant Contents

##### Vitamin C (VC) Content

VC content was obtained by titration with 2,6-dichlorophenol indophenol (DCIP) [26]. Vitamin C content was calculated from the consumption of the DCIP solution.

##### Total Phenols Content (TPC)

We mixed 0.5 g of sample powder with 10 mL anhydrous methanol extractant, which was shaken evenly. After 30 min of ultrasonic extraction at 50 °C and centrifuging at 5 000 ×*g* for 15 min, the supernatant was extracted twice with the same extractant at 10 mL. The residue was washed with a small amount of anhydrous methanol and combined with the supernatant. The supernatant was stored at −40 °C for the TPC and TFC assays.

The TPC was assayed following the Folin–Ciocalteu method reported by Goulas et al. [27] with a few modifications. The extraction solution (0.5 mL) was mixed with FC reagent (0.5 mL) and distilled water (5 mL). After standing for 3 min, 1 mL of sodium carbonate solution (10% *m/v*) was added, and the absorbance at a wavelength of 725 nm was measured after reaction in light for 1 h. Gallic acid was used as the standard sample to create the standard curve, and TPC is expressed as mg/g.

##### Total Flavonoids content (TFC)

The aluminum nitrate colorimetric method by Wang et al [28] was applied to determine the TPC, with certain alterations. About 0.3 mL of sodium nitrite (5% *m/v*) was added to the extraction solution (2.4 mL). After blending and reacting for 6 min, 0.3 mL of aluminum nitrate (10% *m/v*) was added. Then, after reacting for another 6 min, 4 mL of 1 M sodium hydroxide was added, and the absorbance at a wavelength of 510 nm was measured after reaction in light for another 15 min. Rutin was used as a standard sample to create the standard curve, and the TFC is expressed as mg/g.

##### Respiratory Intensity (RI)

The RI was measured using a GXH-3051H infrared carbon dioxide fruit and vegetable respiration tester (Jun-Fang-Li-Hua Technology-Research Institute Beijing, China) in units of mg·CO_2_·kg^−1^·h^−1^. We selected randomly 15 fruits for measurement after weighing.

##### Malondialdehyde (MDA)

MDA content (mmol/g) was measured using the thiobarbituric acid (TBA) method described by Dhindsa et al. [29]. We ground 1.0 g peel to a fine powder and then homogenized it in 10 mL of 50 mM phosphate buffer saline (PBS, pH 7.8). The homogenate was centrifuged at 12,000 ×*g* for 20 min at 4 °C and 2 mL of the supernatant was mixed with 2 mL of TBA, heated at 100 °C for 30 min, and then rapidly cooled. The supernatant was collected by centrifugation at 4 °C and 3000 ×*g* for 20 min at 4 °C. Absorbance was measured at 450, 532, and 600 nm. MDA content was calculated as follows: (mmol/g) = (6.45 × (A_532_-A_600_) - 0.559 × A_450_).

##### Relative Electrical Conductivity (REC)

For relative conductivity, 2.0 g of peel was removed with a 1 cm diameter puncher, rinsed twice in distilled water, and blotted dry with clean filter paper. We then added 20 mL distilled water to the cup and immersed the surface of the peel. After 20 minutes, a conductivity meter (DDS-307A; Rex Shanghai, China) was used to measure the initial conductivity (P_0_) of the sample, and then the sample was boiled for 10 minutes to completely kill the tissue. Then, the sample was cooled to room temperature, and the final conductivity (P_1_) was assayed; relative electrical conductivity = P_0_/P_1_ × 100%.

##### Hydrogen Peroxide (H_2_O_2_)

The H_2_O_2_ content was assayed using the molybdate colorimetric method described by a H_2_O_2_ test kit built in Nanjing (Nanjing Jiancheng Bioengineering Institute, Nanjing, China). We mixed 0.5 g peel powder after grinding with a sample machine with 9 × volume of saline according to the ratio of weight (g):volume (mL) = 1:9. The supernatant was centrifuged for 10 min at 10,000 ×*g*. For the remaining steps, we followed the kit instructions.

The H_2_O_2_ content was calculated as follows: (mmol/g) = (determined OD value - blank OD value)/(standard OD value - blank OD value) × standard concentration (163 mmol/L)/sample weight to be tested.

#### Determination of Antioxidant Enzyme Activities

##### Peroxidase (POD; EC 1.11.1.7) Activity

The POD activity was assayed using the guaiacol method reported by Jin et al. [30]. We extracted 0.5 g of frozen peel with 5 mL of extraction buffer (containing 1% Triton X-100, 1 mmol polyethylene glycol (PEG), 4% polyvinylpyrrolidone) and then centrifuged at 12,000 ×*g* for 30 min at 4 °C. The supernatant obtained by centrifugation is the crude extract of the enzyme. We mixed 50 μL of the extraction with 200 μL of 0.5 mM H_2_O_2_ and 3.0 mL, 25 mM of guaiacol. The OD value of the reaction system was recorded at 470 nm every 10 seconds. More than 6 data records were continuously measured.

##### Superoxide-Dismutase (SOD; EC 1.15.1.1) Activity

The SOD activity was measured using the Nanjing Superoxide Dismutase Test Kit (Nanjing Jiancheng Bioengineering Institute, Nanjing, China). We extracted 0.5 g of frozen peel with 5 mL of extraction buffer and then centrifuged at 12,000 ×*g* for 20 min at 4 °C. The supernatant obtained by centrifugation is the crude extract of the enzyme. Other steps were conducted according to the manufacturer’s instructions. When the inhibition rate of SOD in 1 mL of tissue reaches 50%, the corresponding SOD amount is one unit of SOD activity (U/g).

##### Catalase (CAT; EC 1.11.1.6) Activity

The CAT activity was determined following the method reported by Havir et al. [31] and Robert et al. [32]. We extracted 0.5 g of peel with 5 mL extraction buffer (containing 5 mM dithiothreitol and 5% polyvinylpyrrolid) and then the extraction was centrifuged at 4 °C at 12,000 ×*g* for 15 minutes. The supernatant obtained by centrifugation is the crude extract of the enzyme. The reaction system for measuring CAT activity consisted of 80 μL of supernatant and 2.9 mL of 20 mM H_2_O_2_. The absorbance value of the reaction system was recorded at 240 nm every 30 seconds. More than 6 data points were continuously measured.

##### Ascorbate Peroxidase (APX; EC1.11.1.11) Activity

The APX activity was measured by spectrophotometry. We mixed 0.5 g of peel with 5 mL of extraction buffer (containing 0.1 mM ethylene diamine tetraacetic acid, 0.5 mM ascorbic acid, and 2% polyvinylpyrrolidone) and then the extraction was centrifuged at 12,000 ×*g* for 30 min at 4 °C. The supernatant obtained by centrifugation is the crude extract of the enzyme. We collected 080 μL of the supernatant and mixed it with 0.3 mL of 2 mM H_2_O_2_ and 2.6 mL of reaction buffer (containing 0.5 mM ascorbic acid and 0.1 mM EDTA).

The enzyme activity of each sample was determined using three replicates, and one unit of activity is defined as 0.01ΔOD_470_ ·g^−1^·min^−1^, 0.01ΔOD_240_ ·g^−1^·^−1^·min^−1^, and 0.01ΔOD_290_ ·g^−1^·min^−1^, for POD, CAT, and APX, respectively.

### Statistical Analysis

Drawings were completed with GraphPad Prism 7 software (Auspep, Parkville, Australia);. The treatment effect was determined using the Duncan new complex range method, and SPSS 22.0 software (IBM SPSS 22.0, Chicago, IL) (significance levels at *p* < 0.05 or *p* < 0.01) was used to calculate all the statistical analyses. Correlations between the indicators were analyzed using the Pearson correlation method. Data are presented as means ± standard deviation (SD) with *n* = 3.

## Results and Discussion

### Effects of Different Storage Temperatures on Decay Rate and Weight Loss

The decay rate and weight loss rate are important indexes for evaluating the storability of citrus fruit [33]. Figure 1A illustrates the decay rate in each group was almost zero during the first 30 days of storage. The rotten fruit appeared in 10, 15, and 20, and 5 °C groups at 45 and 75 days, respectively. The decay rate of fruits stored at low temperature was 3.75-fold lower at the end of storage than that of fruits stored at 20 °C. We found significant differences in each treatment group (*p* < 0.01). Numerous previous studies reported that low temperature storage slows the decay of citrus [3,34]. Low temperature storage may reduce the occurrence of fruit diseases and insect pests [12]. In line with our findings, low temperature storage reduced the decay rate of Ponkan mandarins.

**Figure 1.**
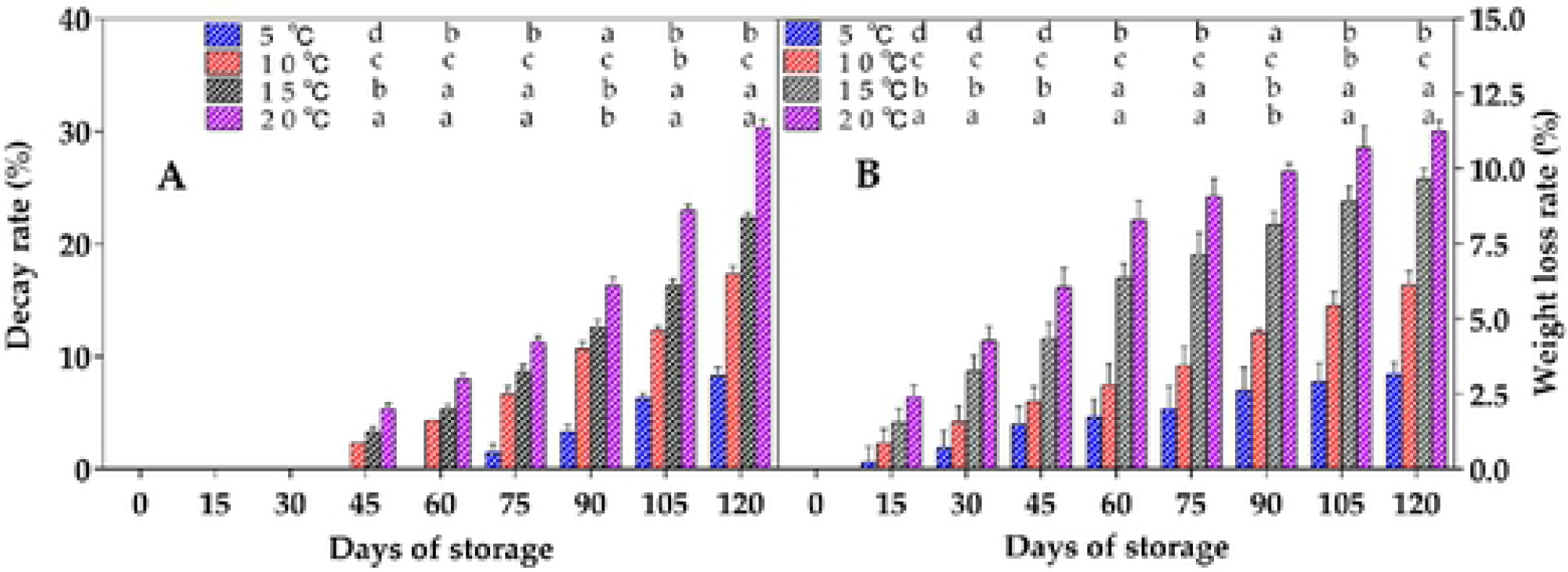
Effect of different storage temperature on (A) decay rate and (B) weight loss of Ponkan mandarins. Data represent the mean ± SD (*n* = 3). Different letters (a, b, c, and d) indicate significant differences (*p* < 0.05) between treatments on the day of sampling.

The weight loss of each group increased with storage time, as shown in Figure 1B, which was mainly caused by water loss due to transpiration. Low temperature storage was shown to significantly reduce the weight loss of Ponkan mandarins (*p* < 0.01). The lowest weight loss was 3% at 5 °C, which is 3.67-fold lower than that of at 20 °C. As some studies reported, low temperature storage can reduce fruit weight loss rate [9,25], which may be due to the slow water transpiration of fruits caused by low temperature, creating a relatively high humidity environment and maintaining the moisture content of fruits.

### Effects of Different Storage Temperatures on CCI, L*, TA, and TSS

As many studies reported, low temperature storage at a suitable temperature is beneficial to maintaining the appearance, color [35], taste, and quality of fruits, improving the storage resistance, and prolonging the shelf life of fruits. To some extent, the color of the fruit reflects the freshness and quality of the fruit. The CCI of Ponkan mandarins under storage at different temperatures increased gradually with the increase in storage time (Figure 2A). However, under low temperature storage, the rate of CCI increase was significantly slower (*p* < 0.05) than at 15 or 20 °C. The value of L^*^ decreased slightly with the prolongation of storage time (Figure 2B). During the whole refrigeration period, L^*^ was lower and significant (*p* < 0.05) at 15 and 20 °C than at 5 and 10 °C under low temperature storage. In this experiment, Ponkan mandarins maintained a good appearance and luster after long storage at a suitable storage temperature. These results are consistent with those reported in other studies [36,37]. Temperature affects the accumulation of carotenoids in citrus fruits, thus significantly affecting its coloration. At the optimum temperature, the more the yellow carotenoids accumulate in the pericarp, the better the color change, the higher the color index, and the brighter the appearance.

**Figure 2.**
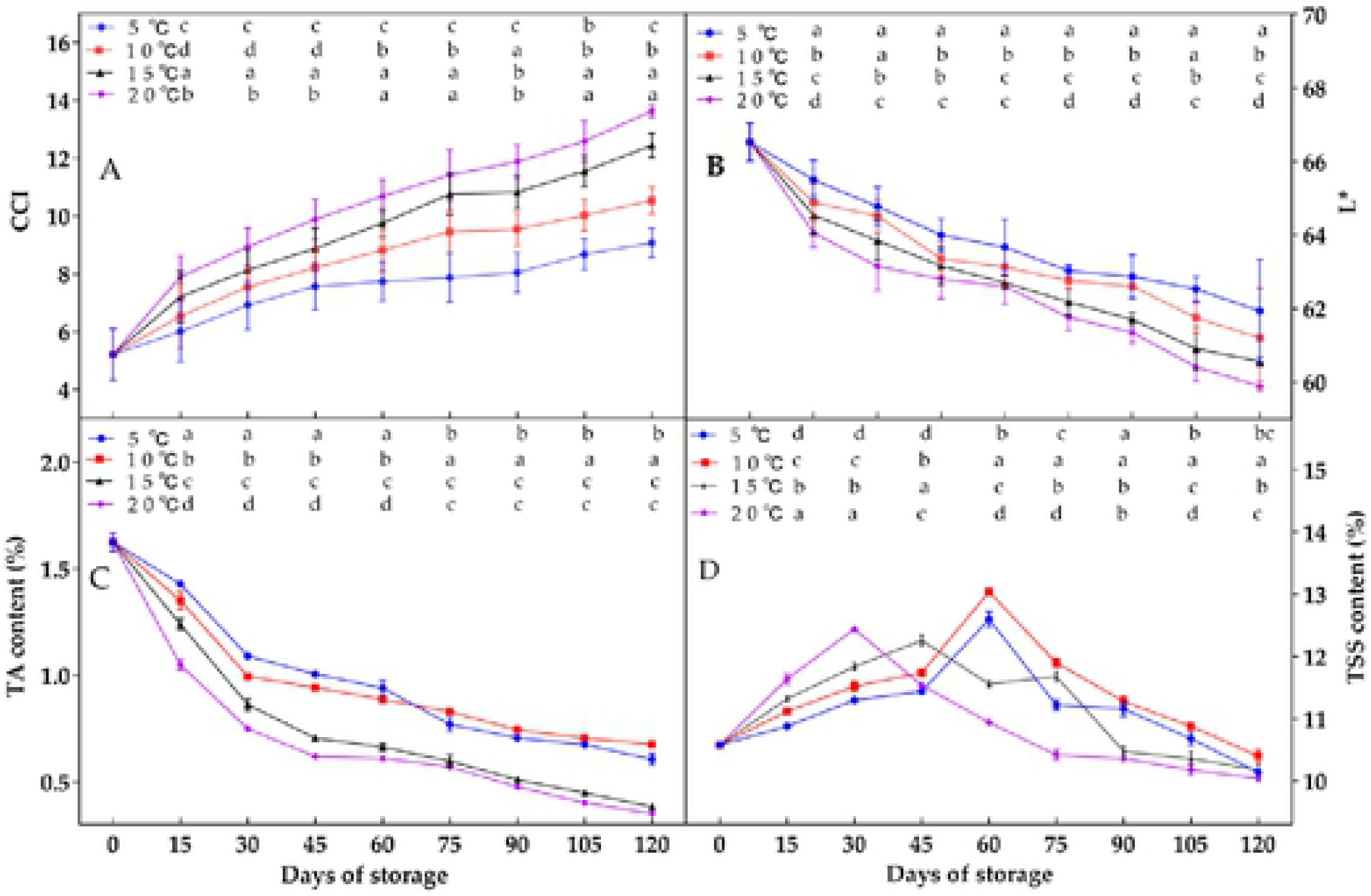
Effect of different storage temperature on (A) CCI, (B) L*, (C) TA content, and (D) TSS content of Ponkan mandarins. Data arc the mean ± SD (*n* = 3). Different letters (a, b, c, and d) indicate significant differences (*p* < 0.05) between treatments on the day of sampling.

As one of the organic acids involved in plant respiration, TA content is regarded as an important index for evaluating the respiration rate of horticultural crops [25]. In our study, after storage for 120 days, the TA content was significantly lower at 10 °C than at 15 and 20 °C (*p* < 0.01) (Figure 2A). The TA content of the fruit at 10 °C retained the initial level of 41.7%, whereas only 21.4% was retained at 20 °C. In this study, the high TA content at 10 °C may be due to the slowed respiration rate, which delayed the degradation of TA [15].

TSS can be maintained at a high level under low temperature storage. TSS in citrus juice is composed of acid soluble pectins, vitamins, sugars, and some soluble proteins [38]. The TSS content of fruits treated at different temperatures increased first and then decreased during the whole storage period. The peaks of TSS content in each treatment group differed, and then the TSS content of each treated fruit decreased rapidly (Figure 2B). The TSS content of fruits at 10 °C was significantly higher (*p* < 0.05) than other treatments (Figure 2B) during the middle and late storage periods (60–120 days). Similar results were reported by Alhassan et al. [15] for Afourer mandarins and Navel oranges.

### Effects of Different Storage Temperatures on Antioxidant Contents

VC is one of the key factors used to evaluate the quality of mandarin fruit. As shown in Figure 3A, the VC content of fruits stored at different temperatures first increased and then decreased during the storage period. The peak times of VC content in each treatment group differed (e.g., 45 days for 10 °C and 30 days for 5, 15, and 20 °C) with the value decreasing 7.63%. The VC content was significantly different from the other treatments in the later storage period (*p* < 0.05). These results are similar to those reported by Zipora et al. [15]; the storage temperature was found to be an important factor affecting the flavor of citrus. Therefore, determining the best minimum safe storage temperature for each citrus variety is important.

**Figure 3.**
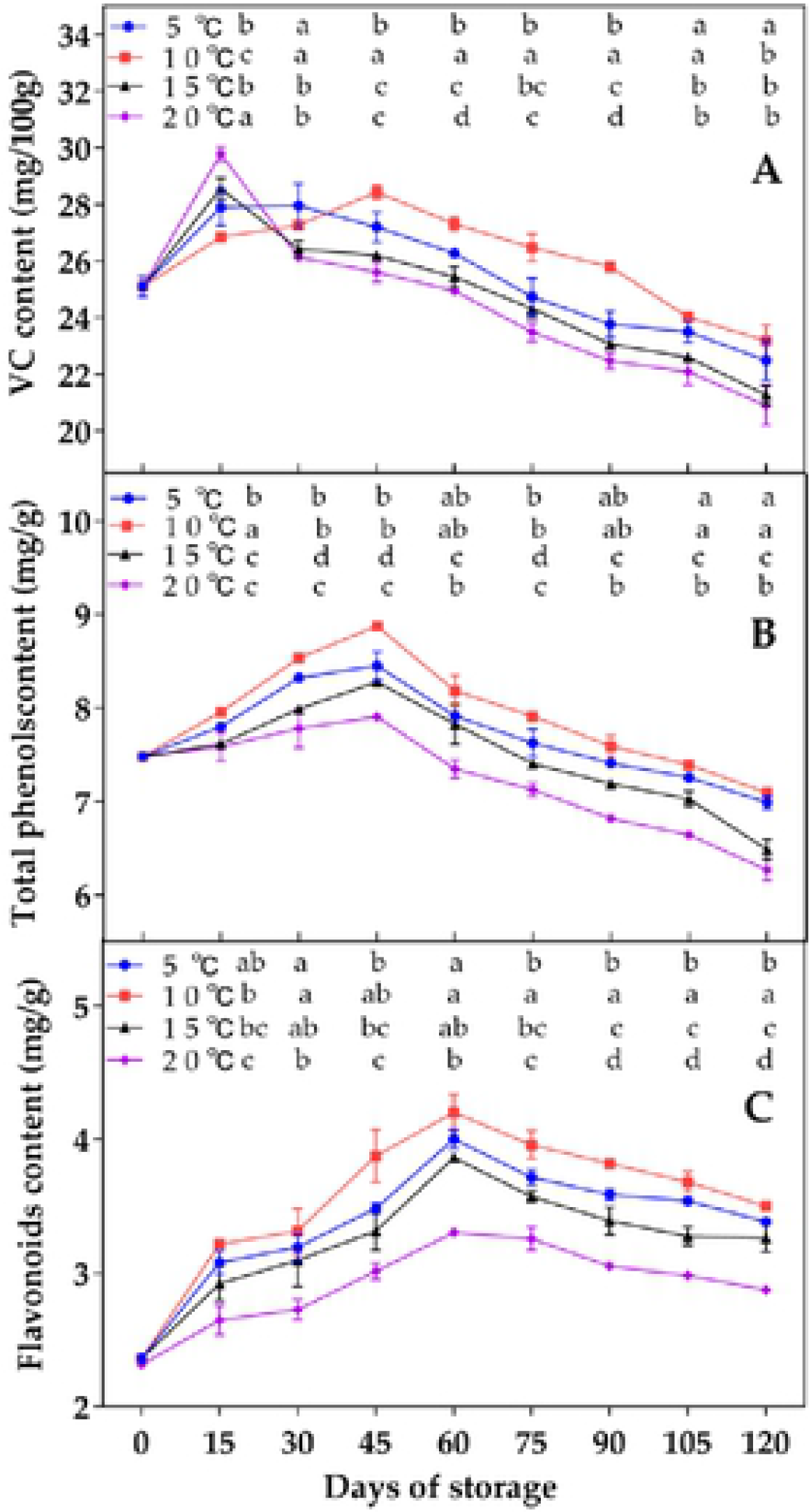
Effect of different storage temperature on (A) VC content, (B) total phenol content TPC, and (C) total flavonoid content TFC of Ponkan mandarins. Data represent the mean ± SD (*n* = 3). Different letters (a, b, c, and d) indicate significant differences (*p* < 0.05) between treatments on the day of sampling.

Phenols and flavonoids are important secondary metabolites in plants. Most of them have the capacity to scavenge free radicals and play an important role in the plant defense mechanism [39]. The contents of total phenols and flavonoids in Ponkan mandarins increased at first and then decreased at different storage temperatures (Figures 3B, C). Total phenol content reached a peak at 45 days and then decreased (Figure 3A). The decrease at 10 °C was slower and the rate was significantly lower than that of the other three treatments (*p* < 0.05). The changes in flavonoids content were similar to that of total phenol content, reaching the peak at 60 days, then decreasing. At 10 °C, the TFC was 1.49 times higher than that at the beginning of storage (Figure 3B), and was significantly (*p* < 0.05) different from that of the other three treatments. In this study, high TPC and TFC were maintained at 10 °C (Figures 3B, C), consistent with previous reports that low temperature storage can maintain high levels of TPC and TFC [40]. In this study, the results indicated that the appropriate storage temperature could improve the disease resistance of fruits by increasing the secondary metabolites with defensive ability in fruit tissues.

### Effects of Different Storage Temperatures on Respiratory Intensity, MDA, REC, and H_2_O_2_

The respiratory intensity of Ponkan mandarins increased under different temperature treatments (Figure 4A). Low temperature storage significantly slowed the increase in the respiratory rate of Ponkan mandarins, and a significant difference (*p* < 0.05) at 5 and 10 °C was found between the earlier and later stage of storage and at 15 and 20 °C (*p* < 0.05). Similar results were reported by Cheng et al. [41] for *Annona chinensis* storage; the respiratory rate of *A. chinensis* storage at 4 °C was significantly lower than that at 8 °C. This is analogous to low temperature storage, which can reduce the respiration of fruits during storage.

**Figure 4.**
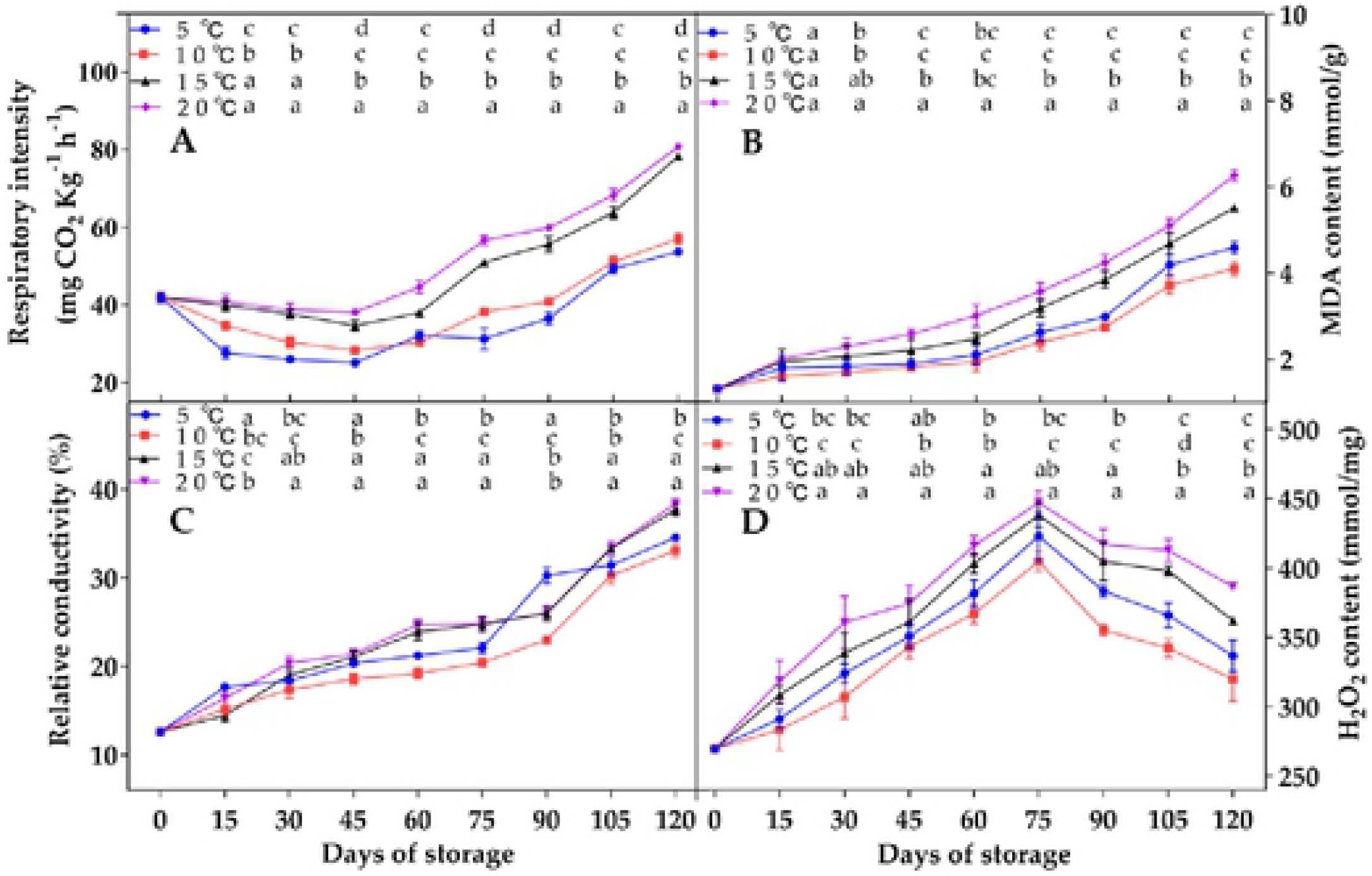
Effect of different storage temperature on (A) respiratory intensity, (B) MDA content, (C) REC, and (D) H_2_O_2_ content of Ponkan mandarins. Data represent the mean ± SD (*n* = 3). Different letters (a, b, c, and d) indicate significant differences (*p* < 0.05) between treatments on the day of sampling.

Active oxygen metabolism increased during storage, reactive oxygen species (ROS) accumulation destroyed cell membrane structure, and MDA content subsequently increased [42,43], which then accelerated fruit senescence. The MDA content at the beginning of fruit storage was 1.3 mmol g^−1^. At the time of storage, the MDA content of the treated group increased, and that of the 10 °C group increased 3.15-fold, which was lower than the other three treatment groups, and was significant (*p* < 0.05) (Figure 4B). MDA is the final product of membrane lipid peroxidation, which is closely related to aging and is one of the direct indicators of membrane oxidative damage. These results are similar to those reported by Wang et al. [16], where low temperature storage at 0 and 10 °C inhibited the increase in MDA content in cherry fruits compared with high temperature storage at 20 and 30 °C. Therefore, the inhibition of MDA content may be related to the senescence and high VC content of fruits stored at low temperature. To further understand the reason for this change, in future analysis, we should determine the protective enzyme activity that delays lipid peroxidation and cell aging.

The relative electrical conductivity of Ponkan mandarins increased at different storage temperatures (Figure 4C) because the gradual senescence of fruits during storage may lead to increased permeability of the pericarp cell membranes. After storage for 120 days, relative conductivity was 30.1% at 10 °C, and 32.6%, 36.2%, and 39.2% for 5, 15, and 20 °C, respectively. The relative conductivity was significantly lower (*p* < 0.05) at 10 °C than for the other three treatments.

Hydrogen peroxide is one of the important representatives of ROS, and its accumulation will cause fruit senescence. During the whole storage process, the H_2_O_2_ content of Ponkan mandarins increased first and then decreased, but increased overall (Figure 4D). At 10 °C, the increase in H_2_O_2_ content was lower compared with the initial storage, by only 1.19 times, significantly lower than the other three treatment groups (*p* < 0.05). In this study, low temperature storage delayed fruit senescence, resulting in a reduction in the accumulation of hydrogen peroxide.

### Effect of Different Storage Temperatures on Antioxidant Enzyme Activities

The activities of POD, CAT, SOD, and APX are closely related to antioxidation and anti-aging in plant tissues. Those enzyme activities in Ponkan mandarins increased at first and then decreased at different storage temperatures (Figure 5).

**Figure 5.**
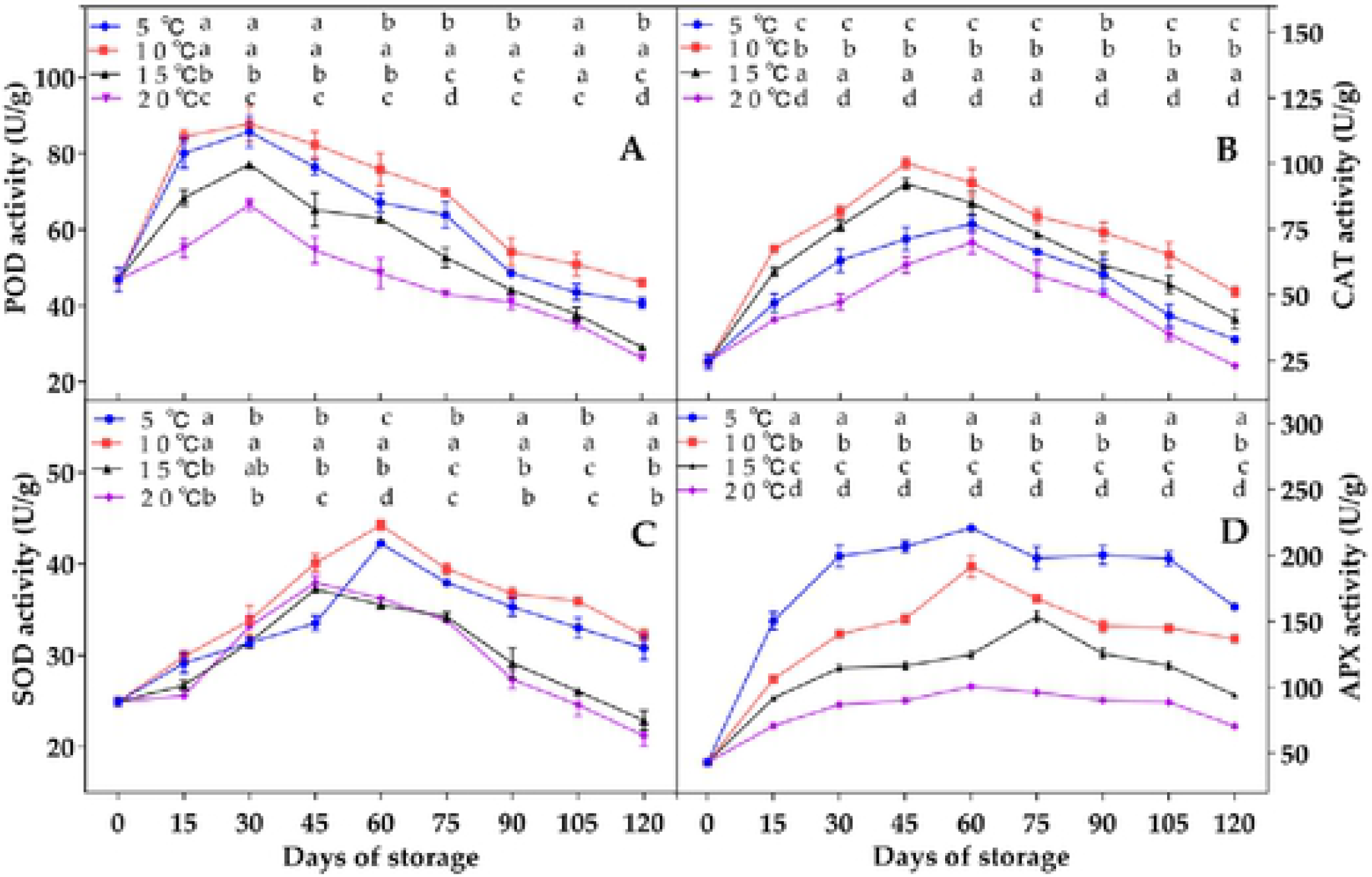
Effect of different storage temperatures on (A) POD, (B) CAT, (D) SOD, and (D) APX activity of Ponkan mandarins. Data represent the mean ± SD (*n* = 3). Different letters (a, b, c, and d) indicate significant differences (*p* < 0.05 or *p* < 0.01) between treatments on the day of sampling.

The POD activity peaked at 30 days and then decreased rapidly (Figure 5A). The rate of decline at 10 °C was significant (*p* < 0.05), which was slower than the other three treatments. The CAT activity was significantly higher (*p* < 0.05; Figure 5B) at the later stage of storage at 10 °C, and the decline rate was slower than that of the other three storage treatments. The SOD activity was 1.29 times higher than that of the initial storage (Figure 5C), and significantly higher than that of the other three treatments (*p* < 0.05). APX activity decreased significantly during storage at 15 and 20 ° C compared with lower temperatures (*p* < 0.05; Figure 5D).

The excessive production and accumulation of ROS caused by fruit senescence destroys the integrity of the cell membrane and reduces the storage tolerance of fruit [42,43]. The high activity of antioxidant and defense-related enzymes can effectively reduce the accumulation of ROS and MDA, reduce oxidative damage, delay fruit aging, and prolong storage life. As important antioxidant enzymes, POD and SOD can prevent membrane lipid peroxidation caused by excessive active oxygen [44,45]. CAT plays an important part in the scavenging process of ROS. As a stress reaction, CAT usually changes with the change in ROS level. APX is the key enzyme for removing hydrogen peroxide in chloroplasts and the main enzyme for vitamin C metabolism. In our study, the results indicated that low temperature storage enhanced the activities of POD, CAT, SOD, and APX, and lessened the accumulation of MDA (Figures 4 and 5). These results are consistent with those of Wang et al. [16], where, compared with high temperature storage at 20 and 30 °C, low temperature storage at 0 and 10°C inhibited the increase in antioxidant enzyme activities in cherry fruits.

### PCA and Pearson Correlation of Different Indexes

Considering the differences in the fruit quality indices of Jing’an Ponkan mandarin under different storage temperatures and according to the results of the previous experiments, 17 indexes of Ponkan mandarins during postharvest storage were standardized using PCA with SPSS 20.0 software (IBM SPSS 22.0, Chicago, IL). Wang et al. [46] compared the differences among different citrus varieties, and then established a citrus quality evaluation system using PCA as the basis for evaluating the comprehensive quality of different citrus juices. PCA [24] was used to further integrate and analyze the results of fruit quality indicators. By analyzing the eigenvalues of the covariance matrix, the first two principal components (PCs) account for 83.19% of the total variance of the data set. PC1 explained 53.622% of the variance in the data set, and PC2 explained 29.568%. PC1 had high positive loading for weightlessness, relative conductivity, MDA content, and CCI, and a high negative loading for TPC, VC content, and POD activity. PC2 had high positive loading for SOD, CAT, and APX activities (Figure 6).

**Figure 6.**
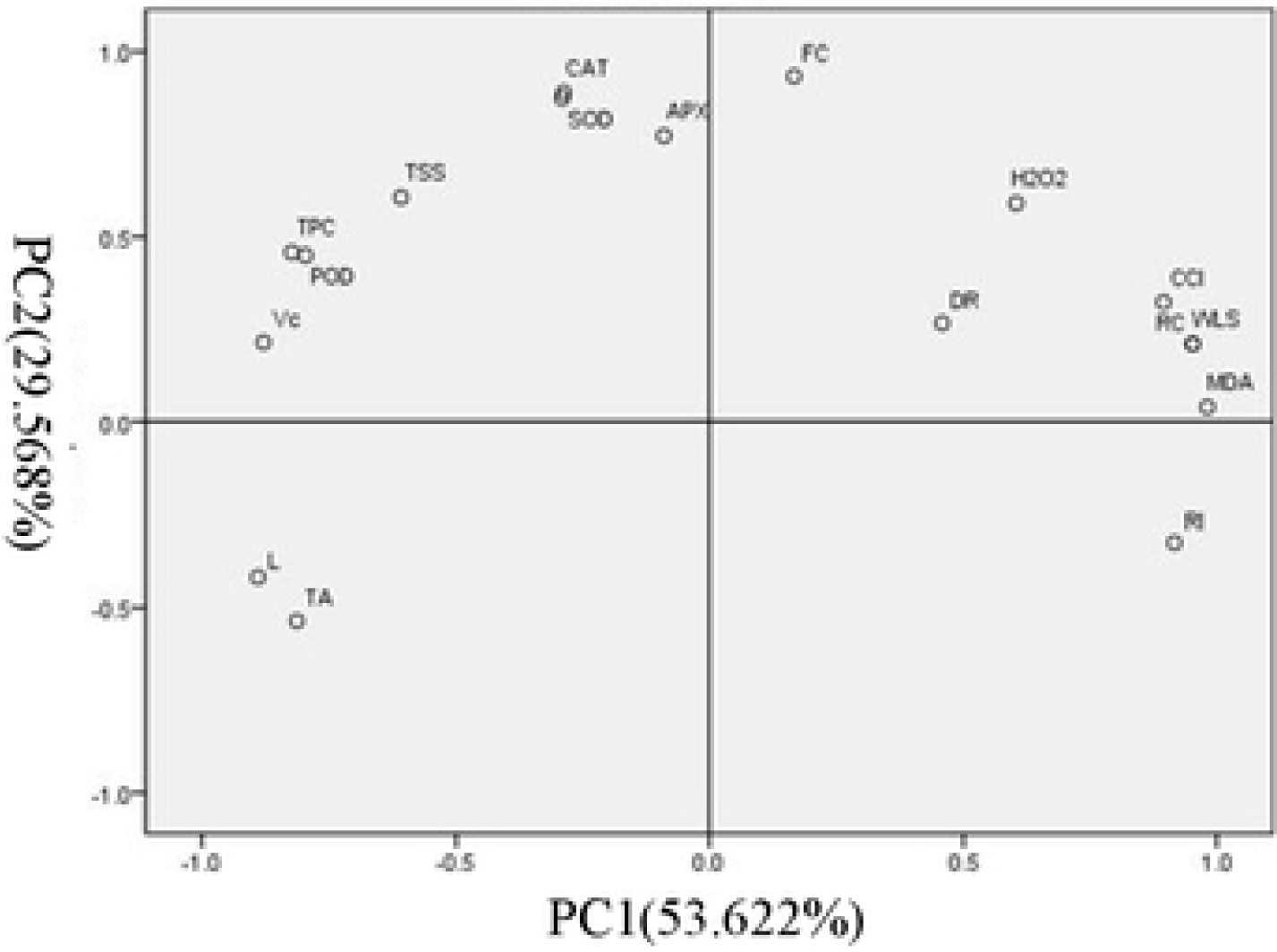
Two-dimensional principal component analysis (PCI: 53.622%, PC2: 29.568%) of various indicators of Ponkan mandarins under different storage temperatures. DR, decay rate; WLR, weight loss rate; TPC, total phenols content; TFC, total flavonoids content; RI, respiration intensity; REC, relative conductivity.

After 60 days of storage, the score dropped sharply (Figure 7), indicating that the optimal storage time of Ponkan mandarins is 60 days. When the storage time is prolonged, the antioxidant and anti-aging ability of the fruit gradually decreases. The comprehensive score of 10 °C storage fruits was always the highest among the four storage temperatures, which maintained good fruit quality. Therefore, the optimum storage temperature of Ponkan mandarins is 10 °C.

**Figure 7.**
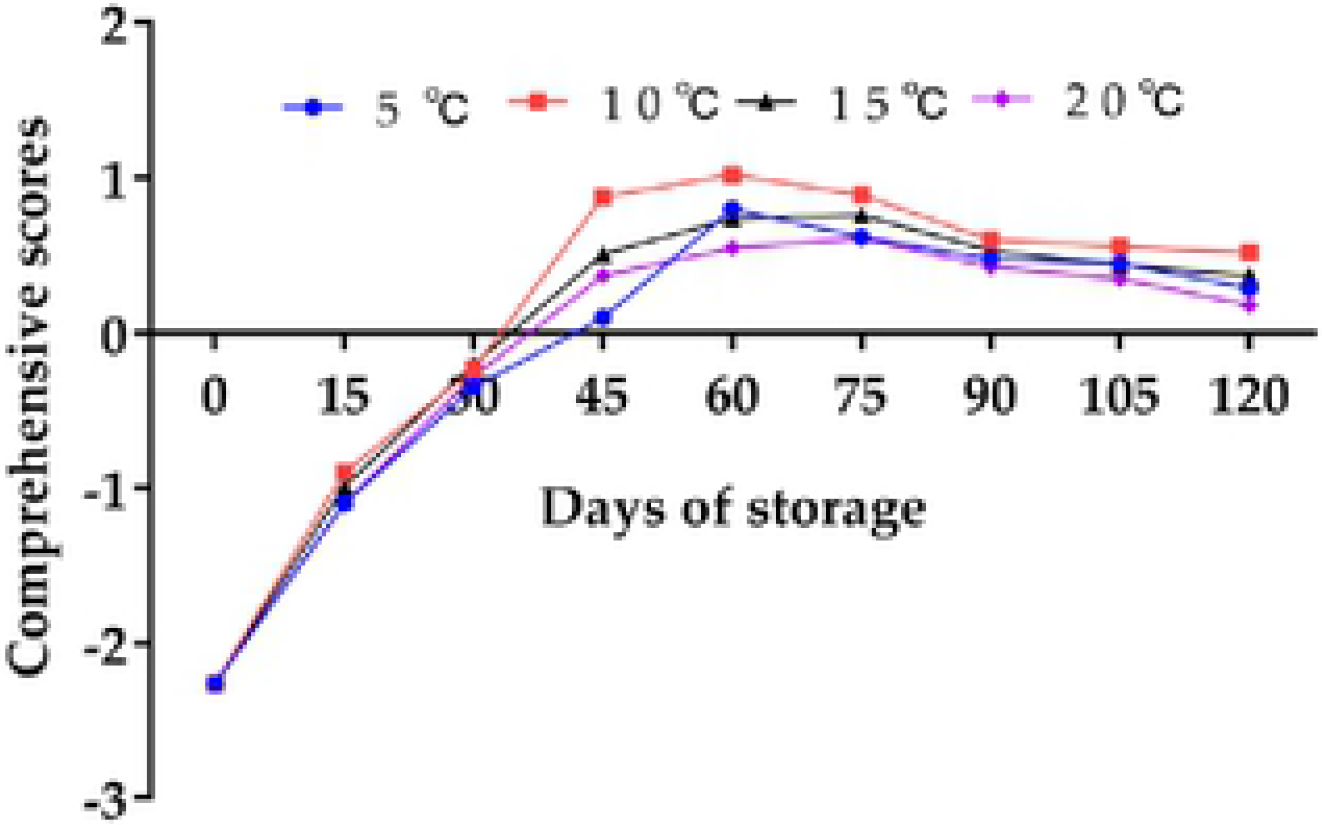
Comprehensive evaluation score among four storage temperatures.

A correlation-based approach using the Pearson coefficient was used to examine the positive and negative relationships between the indexes of Ponkan mandarins at different storage temperatures. Significant positive correlations (in red) and negative correlations (in blue) are displayed in Figure 8. Notably, VC content was significant negatively correlated with weight loss (*r* = 0.791, *p* < 0.01, CCI (*r* = 0.647, *p* < 0.01)), respiratory intensity (*r* = 0.852, *p* < 0.01), and MDA content (*r* = 0.837, *p* < 0.01). However, VC content was significantly positively correlated with TSS content (*r* = 0.708, *p* < 0.01), TPC (*r* = 0.838, *p* < 0.01), and POD activity (*r* = 0.848, *p* < 0.01). These results indicate that fruit quality is maintained by some antioxidants.

**Figure 8.**
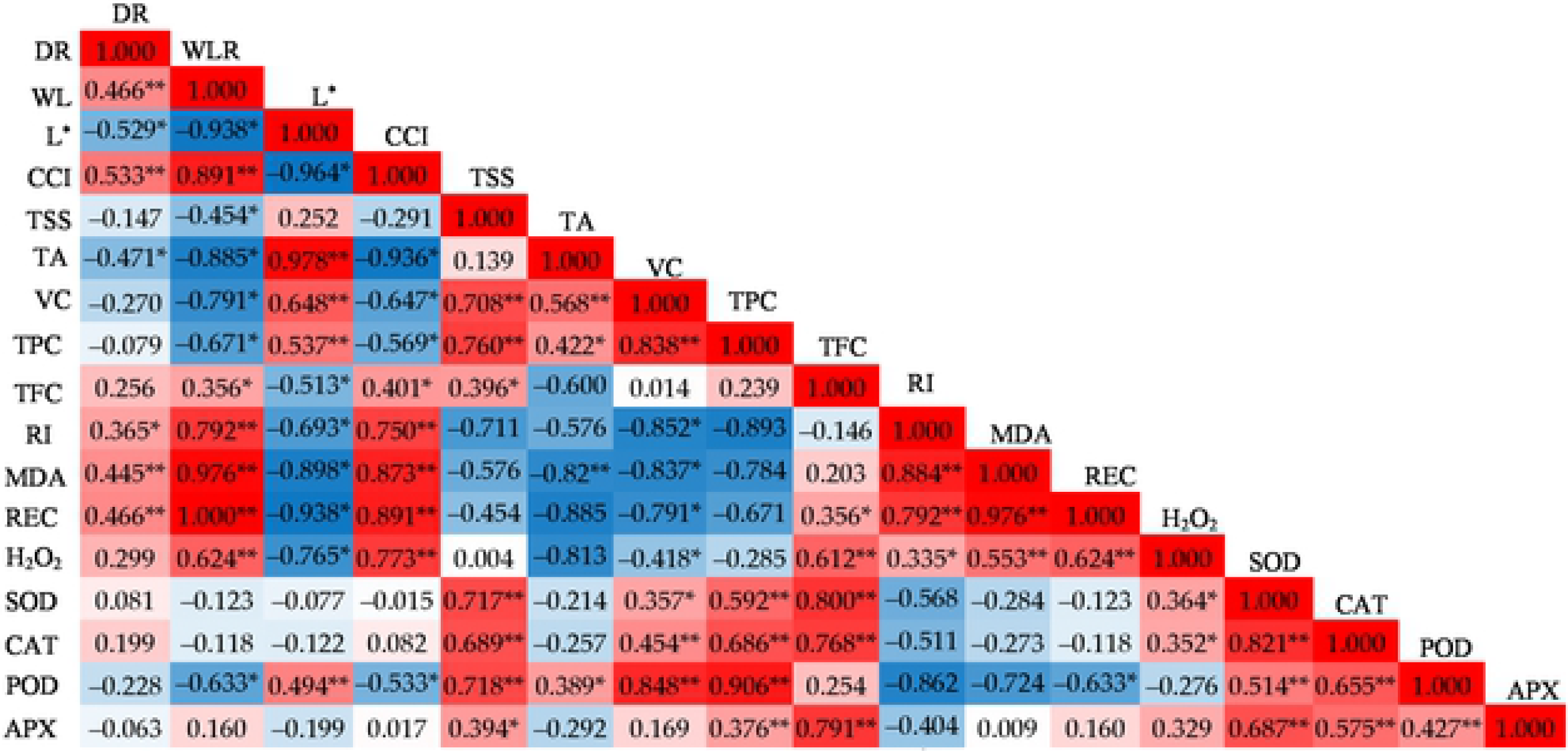
Correlation matrix based on Pearson’s correlation coefficient between the indexes under different storage temperatures was analyzed. The deepness and value in the red and blue grids arc expressed the levels of positive and negative correlation, respectively. The symbol * and ** represented significant difference at *p* < 0.05 and *p* < 0.01, respectively.

MDA content was significantly positively correlated with decay rate (*r* = 0.445, *p* < 0.01), respiratory intensity (*r* = 0.884, *p* < 0.01), respiratory intensity (*r* = 0.976, *p* < 0.01) and H_2_O_2_ content (*r* = 0.553, *p* < 0.01). However, MDA content was negatively correlated with TPC (*r* = 0.784, *p* < 0.01), and POD activity (*r* = 0.724, *p* < 0.01). These results indicate that antioxidants and enzymes scavenge ROS and delayed fruit senescence during storage.

We found a significantly positive correlation between H_2_O_2_ content and SOD activity (*r* = 0.364, *p* < 0.01) and CAT activity (*r* = 0.352, *p* < 0.01). These results indicate that H_2_O_2_ induces the increase in SOD and CAT antioxidant enzymes and scavenges ROS. This result has been confirmed: the accumulation of H_2_O_2_ induced the increase in POD, SOD, CAT, and APX activities [44], which scavenged ROS [37,38], reduced MDA and H_2_O_2_ content in cells, and maintained redox balance [49].

## 4. Conclusions

In conclusion, the storage temperature of 10 °C can effectively maintain the quality of the fruit by improving the storability of Ponkan mandarins, reducing the accumulation of MDA and H_2_O_2_, reducing the level of ROS, maintaining high levels of defense enzyme activities, improving disease resistance, delaying fruit aging, and prolonging shelf life. We did not delve into the molecular mechanism of how low temperature maintains fruit quality, which requires further exploration.

## Author Contributions

conceptualization, J.C. and C.C.; methodology, N.C.; validation, N.C., C.W., and C.C.; formal analysis, N.C.; investigation, N.C.; resources, J.C.; data curation, C.C. and C.W.; writing–original draft preparation, N.C.; writing–review and editing, C.C. and C.W.; supervision, J.C.; project administration, J.C.; funding acquisition, J.C.

